# Whole-brain mapping of monosynaptic inputs to midbrain cholinergic neurons

**DOI:** 10.1101/2020.08.14.251546

**Authors:** Icnelia Huerta-Ocampo, Daniel Dautan, Nadine K. Gut, Bakhtawer Khan, Juan Mena-Segovia

## Abstract

The cholinergic midbrain is involved in a wide range of motor and cognitive processes. Cholinergic neurons of the pedunculopontine (PPN) and laterodorsal tegmental nucleus (LDT) send long-ranging axonal projections that target sensorimotor and limbic areas in the thalamus, the dopaminergic midbrain and the striatal complex following a topographical gradient, where they influence a range of functions including attention, reinforcement learning and action-selection. Nevertheless, a comprehensive examination of the afferents to PPN and LDT cholinergic neurons is still lacking, partly due to the neurochemical heterogeneity of this region. Here we characterize the whole-brain input connectome to cholinergic neurons across distinct functional domains (i.e. PPN vs LDT) using conditional transsynaptic retrograde labeling in ChAT::Cre male and female rats. The quantification of input neurons revealed that both PPN and LDT receive similar substantial inputs from the superior colliculus and the output of the basal ganglia (i.e. substantia nigra pars reticulata). In addition, we found that PPN cholinergic neurons receive preferential inputs from basal ganglia structures than from the cortex, whereas LDT cholinergic neurons receive preferential inputs from cortical areas. Our results provide the first characterization of inputs to PPN and LDT cholinergic neurons. The differences in afferents to each cholinergic structure support their differential roles in behavior.

**Significance statement:** Acetylcholine is a widespread neuromodulator that regulates a wide variety of functions including learning, goal-directed behavior and execution of movements. In this study we characterized the distribution of presynaptic neurons that modulate the activity of functionally distinct midbrain cholinergic neurons located in the pedunculopontine nucleus (PPN) and the laterodorsal tegmental nucleus (LDT) by using a transsynaptic, modified-rabies virus strategy. We reveal that input neurons are widely distributed throughout the brain but segregated into specific functional domains. Motor related areas innervate preferentially the PPN, whereas limbic related areas preferentially innervate the LDT. Our results suggest that input neurons located along distinct functional domains have differential impact over cholinergic midbrain regions.

## Introduction

Acetylcholine is a major neuromodulator that plays a central role in attention, movement and behavioral flexibility. One of the major sources of acetylcholine is located in the midbrain, where cholinergic neurons of the pedunculopontine nucleus (PPN) and laterodorsal tegmental nucleus (LDT) provide widespread innervation to the thalamus (Steriade et al., 1988; Parent and Descarries, 2008; Huerta-Ocampo et al., 2020) and the basal ganglia (Woolf and Butcher, 1986; Clarke et al., 1987; Bolam et al., 1991; Dautan et al., 2014, 2016a). Recent studies using genetic approaches to selectively manipulate the activity of cholinergic neurons have shed light into the functions of cholinergic neurons of the PPN and LDT. For example, cholinergic neurons have been shown to be involved in reinforcement learning through the modulation of dopamine neurons in the ventral tegmental area (VTA) (Dautan et al., 2016b) and induce movement through dopamine-mediated mechanisms (Dautan et al., 2016b; Xiao et al., 2016). In contrast to the classic notions of their involvement in motor activity and wakefulness regulation, however, it has been shown that optogenetic activation of cholinergic neurons in resting mice does not evoke a motor response (Roseberry et al., 2016), and chemogenetic experiments have shown that activation of cholinergic neurons does not increase the amount of time spent in wakefulness (Kroeger et al., 2017). These experiments thus highlight the need to revisit some of the theories of the midbrain cholinergic function. A recent study, for example, has shown that midbrain cholinergic neurons innervate the striatal complex and make direct connections with striatal cholinergic interneurons (CINs); manipulations of either midbrain cholinergic neurons (i.e. PPN/LDT) or CINs have a similar influence on action strategy encoding, suggesting convergent functional roles between midbrain and striatal cholinergic systems (Dautan et al., 2020). Thus, to determine the functional overlap between cholinergic cell groups it is critical to understand how their activity is regulated by their afferent systems. Recent studies have described the input connectivity of cholinergic neurons of the basal forebrain (Do et al., 2016; Gielow and Zaborszky, 2017) and the striatum (Guo et al., 2015; Klug et al., 2018) but the sources of inputs to the cholinergic neurons of the PPN and LDT are still missing.

Cholinergic neurons in the PPN and LDT form a continuum that extends from the caudal end of the substantia nigra (SN) to the central gray matter near the fourth ventricle. The efferent connectivity of these two structures is characterized by a topographical arrangement where PPN neurons preferentially innervate motor neuronal systems, whereas LDT neurons preferentially innervate limbic neuronal systems (Mena-Segovia, 2016). For example, cholinergic neurons of the PPN innervate the dopaminergic neurons of the substantia nigra pars compacta (SNc), the dorsal striatum and thalamic relay nuclei. In contrast, cholinergic neurons of the LDT innervate the ventral striatum and limbic thalamic nuclei. While not entirely segregated, both structures provide converging innervation to a certain subset of structures (i.e. the intralaminar thalamic nuclei, VTA), although detailed examination of the postsynaptic targets at the level of the VTA has revealed a divergent modulation over dopamine efferent circuits (Dautan et al., 2016b), thus suggesting a largely unexplored level of functional selectivity between PPN and LDT cellular targets. Thus, the identification of the afferent systems to PPN and LDT cholinergic neurons is critical to integrate an input/output connectivity map and to understand how their activity is regulated.

Here we aimed to identify and map in whole-brain sections the distribution of presynaptic neurons that specifically synapse onto cholinergic neurons of the PPN and LDT by using a transsynaptic retrograde labeling approach. Our results show a substantial degree of overlap in the structures that innervate both regions but critically reveal a topographical segregation along functionally specialized regions of the cholinergic midbrain that supports previous anatomical and behavioral findings.

## Methods

### Animals

All experimental procedures were performed on adult male and female ChAT::Cre+ rats (Witten et al., 2011). Rats were maintained on a 12 h light/dark cycle (light on 7:00 A.M.) and *ad libitum* access to water and food. All procedures were performed in accordance with the Society for Neuroscience policy on the use of animals in neuroscience and were approved by the Institutional Animal Care and Use Committee of Rutgers University, in compliance with the National Institutes of Health Guide for the Care and Use of Laboratory Animals.

### Stereotaxic Injections

Surgeries were performed under deep isoflurane anesthesia (2% in O2; Isoflo; Schering-Plough). To demonstrate monosynaptic inputs to the PPN and LDT we used a transsynaptic tracing system based on the modified rabies virus strategy (Wickersham et al., 2007; Watabe-Uchida et al., 2012). For this, animals were injected with 500nL of a 1:1 mixture containing rAAV5/EF1a-Flex-TVA-mCherry, titer 4.3e12 VP/mL, and rAAV5/CA-Flex-RG, titer 2e12 VP/mL (both from University of North Carolina vector core). Injections targeted the rostral (500 nl over 10 min; from bregma in mm: AP, −7.3; ML, +1.8; DV, −6.8 ventral of the dura; *n* = 3) and caudal parts of the PPN (500 nl over 10 min; from bregma in mm: AP, −7.8; ML, +1.8; DV, −6.5 ventral of the dura; *n* = 3). However, the results in terms of the distribution of input neurons did not show major differences between these two regions of the PPN and for this reason the data were pooled. Injections were also made in the LDT (500 nl over 10 min; from bregma in mm: AP, −8.5; ML, +0.9; DV, −6.0 ventral of the dura; *n* = 3) (Paxinos and Charles Watson, 2007). All injections were made using designated 1-µl Hamilton syringes at a rate of 50 nl/min and post-injection diffusion time of 5 min. Fourteen days later, 500nL of EnvA-ΔG-rabies-eGFP (Salk vector core, titer 4.3e8 transducing units [TU/mL]) (Wickersham et al., 2007) were injected into the PPN and LDT using the same coordinates, the same rate of injection and diffusion time. Control experiments were performed where the modified rabies was injected to ChAT::Cre rats that did not receive any helper viruses.

### Immunohistochemistry and imaging

Seven days after rabies virus injections, the rats were transcardially perfused with 0.05M PBS, pH 7.4, followed by 300 ml of 4% w/v paraformaldehyde in phosphate buffer (0.1M, pH 7.4). Brains were stored in phosphate buffer saline (PBS) with 0.05% azide at 4°C until sectioning. Sagittal sections of 50 μm thickness were obtained and collected in PBS, using a vibratome (VT1000S; Leica) and organized in series. For each brain, the site of injection was verified and only those with on-target injections were processed further. All the incubations were done in “Triton-PBS” (PBS containing 0.3% v/v Triton X-100 [Sigma]). Every fourth section was blocked for 2h at room temperature (RT) while shaking in Triton-PBS containing 10% v/v of normal donkey serum (NDS; Jackson Immunoresearch). Next, they were incubated overnight in an anti-GFP antibody coupled with a 488 fluorophore (1:1000, Invitrogen, A-21311). Sections containing the sites of injection were incubated in rat-raised anti-GFP (1:1000, Nacalai tesque, 04404-84), rabbit anti-mCherry (1:1000, Abcam, ab167453) and goat anti-ChAT (1:500, Millipore AB 144P) overnight at RT. After washing, the sections were incubated for 4-6 h in the following secondary antibodies: donkey anti-rat 488 (1: 500; Jackson Immunoresearch, 712-546-153), donkey anti-rabbit-Cy3 (1:500; Jackson Immunoresearch,711-165-152) and donkey anti-goat 405 (1:500; Jackson Immunoresearch,711-475-152) in Triton-PBS containing 1% of NDS. ChAT immunostaining was performed to delineate the borders of the PPN and LDT and assess the accuracy of our injections.

To determine the location and neurochemical identity of input neurons localized within the SN and dorsal raphe (DR), selected sections were incubated using antibodies against mouse tyrosine hydroxylase (TH) (1:1000, Sigma, T2928) and rabbit tryptophan hydroxylase 2 (1:500, Novus Biological, NB100-7455), respectively. Sections were then incubated in either donkey anti-mouse 405 (1:500; Jackson Immunoresearch, 715-475-151) or donkey anti-rabbit 405 (1:500; Jackson Immunoresearch, 711-475-152). After several washes, the fluorescently labeled sections were mounted on glass slides using Vectashield and examined on a confocal (FV-2000; Olympus) microscope. Whole-brain images were obtained by tiling and stitching individual images taken at 10x magnification. Sections were scanned at 10, 20 and 30 µm in depth, thus generating three images per brain section.

### Analysis of input neurons

The brightness and contrast of the captured images were adjusted in Photoshop (Adobe Systems) and superimposed with templates modified from an atlas to perform the mapping in the distinct brain areas (Paxinos and Charles Watson, 2007). To quantify input cell numbers, we manually counted eGFP-positive neurons found on each of the three depths of the captured images per whole-brain section. We only recorded a positive neuron when we were able to identify its cell body in its entirety. Sections where only dendritic processes and axons or fractions of cell bodies were identified, were not counted. Thus, it is likely that our counting underestimated the total number of input neurons. We counted and registered in an excel spreadsheet the number and location of every input neuron found in each brain region throughout the most lateral to most medial levels using the counter cell in-built function of the ImageJ software. This was done for all brains included in this study (n=9). Brains with less than a total of 500 input neurons were not considered for analysis. From all the brain nuclei where we found input neurons within the PPN and LDT groups, we selected only those regions in which at least one of 3 animals had more than 5 neurons and the other two animals had more than 1 neuron. If one of the animals did not have any neuron in a given structure, that structure was not included in our final analysis. By applying these criteria, we obtained a list of 50 structures which are reported in this study (see **Table 1** for abbreviations). Cells were counted by two experimenters, one of which remained blind for the entire phase of analysis.

**Table 1.** List of abbreviations.

|  |  |  |  |
| --- | --- | --- | --- |
| **3N** | Oculomotor n. | **NA7** | Nucleus adrenergic 7 |
| **5N** | Motor trigeminal n. | **OFC** | Orbitofrontal cortex |
| **7N** | Facial n. | **PAG** | Periaqueductal gray |
| **ACC** | Anterior cingular cortex | **PC** | Parietal cortex |
| **APT** | Anterior Pretectal nucleus | **PCRt** | Parvicellular reticular n. |
| **BIC** | Brachium of superior colliculus | **PMnR** | Paramedian raphe nucleus |
| **CnF** | Cuneiform n. | **PnC** | Pontine reticular n., caudal |
| **DMTg** | Dorsomedial tegmental area | **PnO** | pontine reticular n., oral |
| **DpG** | Deep gray layer SC | **PP** | peripeduncular n. |
| **DpWh** | Deep white layer SC | **PPN** | Pedunculo pontine n. |
| **DR** | Dorsal raphe | **Pr5** | Principia sen. trigeminal n. |
| **dtgx** | Dorsal tegmental decussation | **PrCnF** | Precuneiform area |
| **Eth** | Ethmoid thalamic n. | **RRF** | Retrorubral Field |
| **Gi** | Gigantocellular reticular n. | **RtTg** | Reticulotegmental n. |
| **GPe** | Globus Pallidus pars externa | **SC** | Superior Colliculus |
| **GPI** | Globus Pallidus pars interna | **SN** | Substantia Nigra |
| **IC** | Inferior Colliculus | **SNc** | SN pars compacta |
| **Int Cajal** | Interstitial nucleus of Cajal | **SS** | Somatosensory cortex |
| **InG** | Intermediate gray layer SC | **STN** | Subthalamic n. |
| **InWh** | Intermediate white layer SC | **Su5** | Supratrigeminal n. |
| **IRtA** | Intermediate reticular n., alpha | **VC** | Visual cortex |
| **LDT** | Laterodorsal n. | **VP** | Ventral Pallidum |
| **MiTg** | Microcellular tegmental n. | **VS** | Ventral striatum |
| **MC** | Motor cortex | **VTA** | Ventral Tegmental Area |
| **mRt** | Mesencephalic reticular form. | **ZI** | Zona Incerta |

### Experimental design and statistical analyses

For the comparison of the distribution of input neurons to PPN and the LDT we compared two experimental groups, rats injected in the PPN for the examination of input neurons to the PPN and rats injected in the LDT for the examination of input neurons to the LDT. For the evaluation of differences between the distribution of input neurons from different functional areas we considered two independent variables (target areas and functional input areas) and conducted 2-way ANOVAs on ln-transformed (to address the violation of the homogeneity of variances) numbers of input neurons followed by univariate tests to understand the simple main effects of the target structure on each functional area. To examine if selected structures within functional areas have more input neurons projecting to the PPN or LDT, respectively, than other structures, we used the Kruskal-Wallis H test, because the variances of the individual groups were heterogeneous. The counts of inputs neurons were normalized to either the total number of input neurons (Fig. 3) or the number of starter neurons (Fig. 4 and 5). Post-hoc tests were Bonferroni-corrected. To compare differences between the number of input neurons to the PPN and LDT from all the individual structures that we identified to have input neurons, we conducted two-sided t-tests.

**Figure 1.**
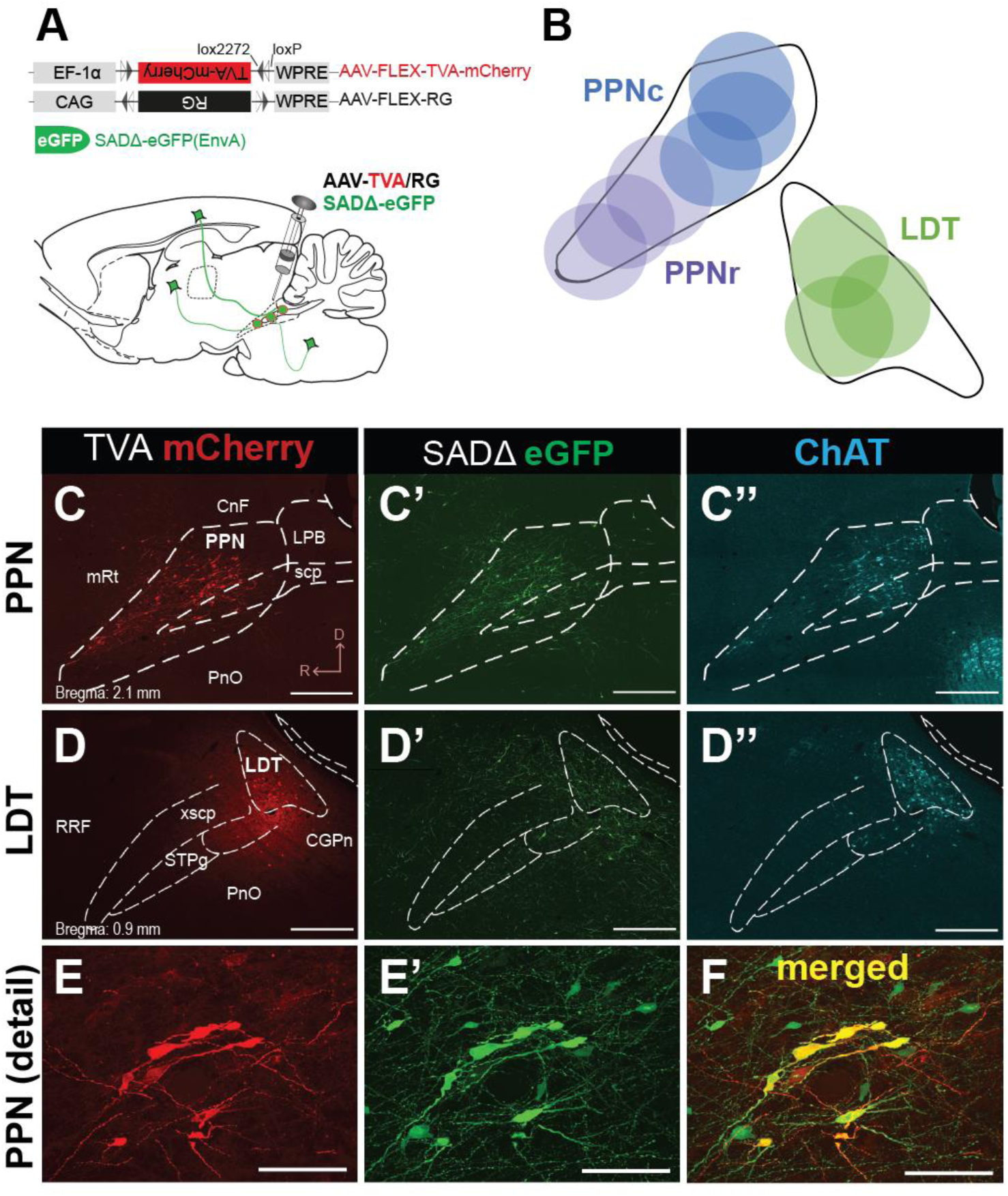
Transsynaptic retrograde tracing of midbrain cholinergic neurons. **A**, Schematic of the experimental procedure. AAV5-FLEX-TVA-mCherry and AAV8-FLEX-RG helper viruses were injected into the PPN or LDT of rats that expressed Cre in cholinergic neurons. Two weeks after these injections, a modified rabies virus SADΔG-eGFP (EnvA) was injected into the same area and the brain was processed after 7 days. **B**, Location of the site of injections in the PPN and LDT. **C-E**, Injections were confined to the borders of the PPN (**C-C’**) and LDT (**D-D’**), as determined by ChAT immunolabeling (**C”** and **D”**). **E**, Starter neurons were identified by the expression of the TVA helper reporter and SADΔ-eGFP. Scale bars C & D: 1 mm. E-F: 100 µm.

**Figure 2.**
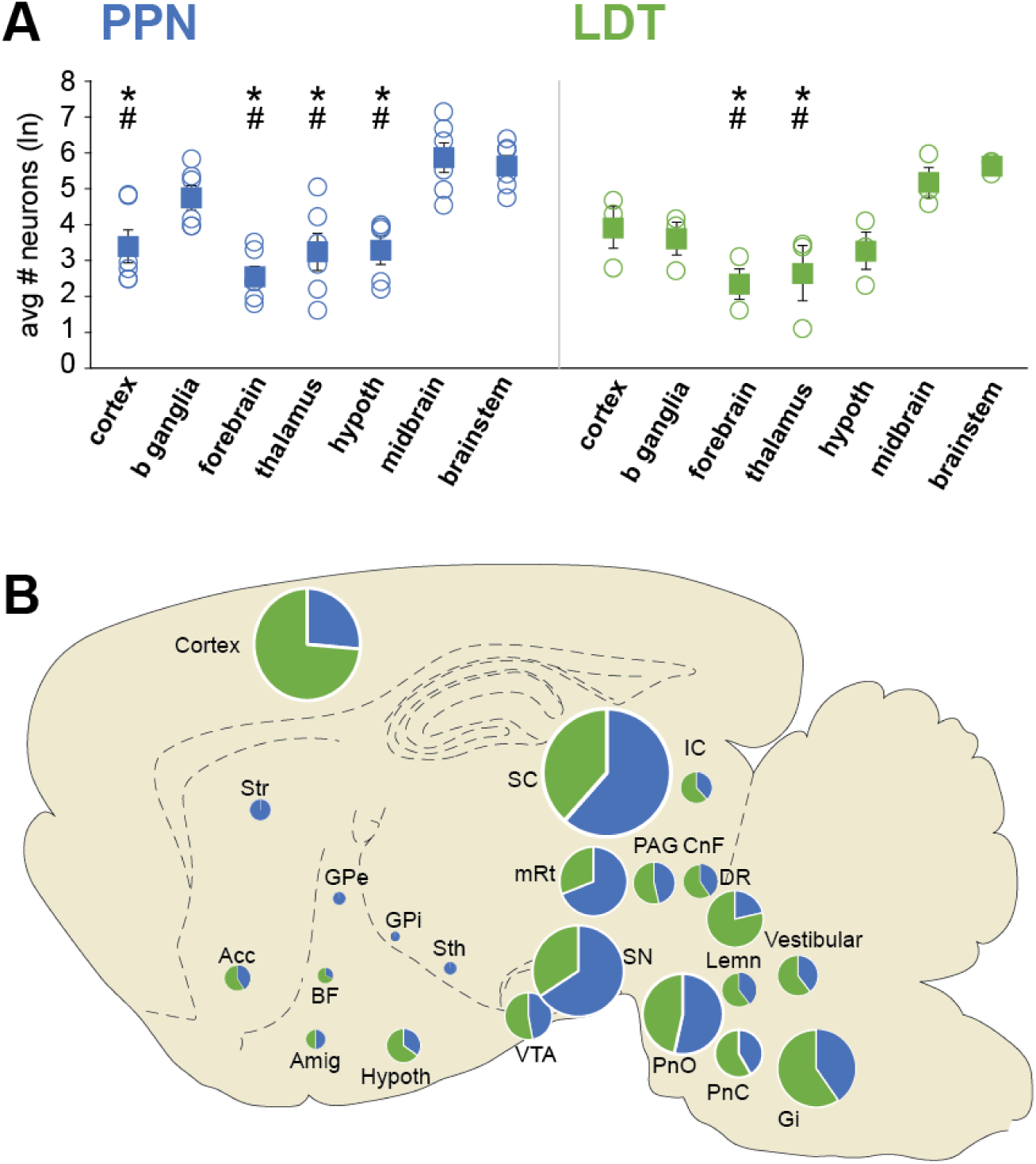
Afferents to PPN and LDT cholinergic neurons are heterogeneously distributed across brain regions. **A**, Average number (mean ± SEM) and individual subject values of input neurons (PPN n=6 rats; LDT n=3 rats) grouped by anatomical region, representing the cortex, basal ganglia, forebrain, thalamus, hypothalamus, midbrain and brainstem. Input neurons were more concentrated in the midbrain and brainstem for both PPN and LDT, and in the basal ganglia for the PPN. *, significant when compared to midbrain; #, significant when compared to brainstem. **B**, Summary schematic illustrating the distribution of input neurons across the brain. The pie charts show the percentage of neurons projecting to the PPN or LDT, whereas the size of the chart shows the relative strength of this connection as estimated by the number of neurons.

**Figure 3.**
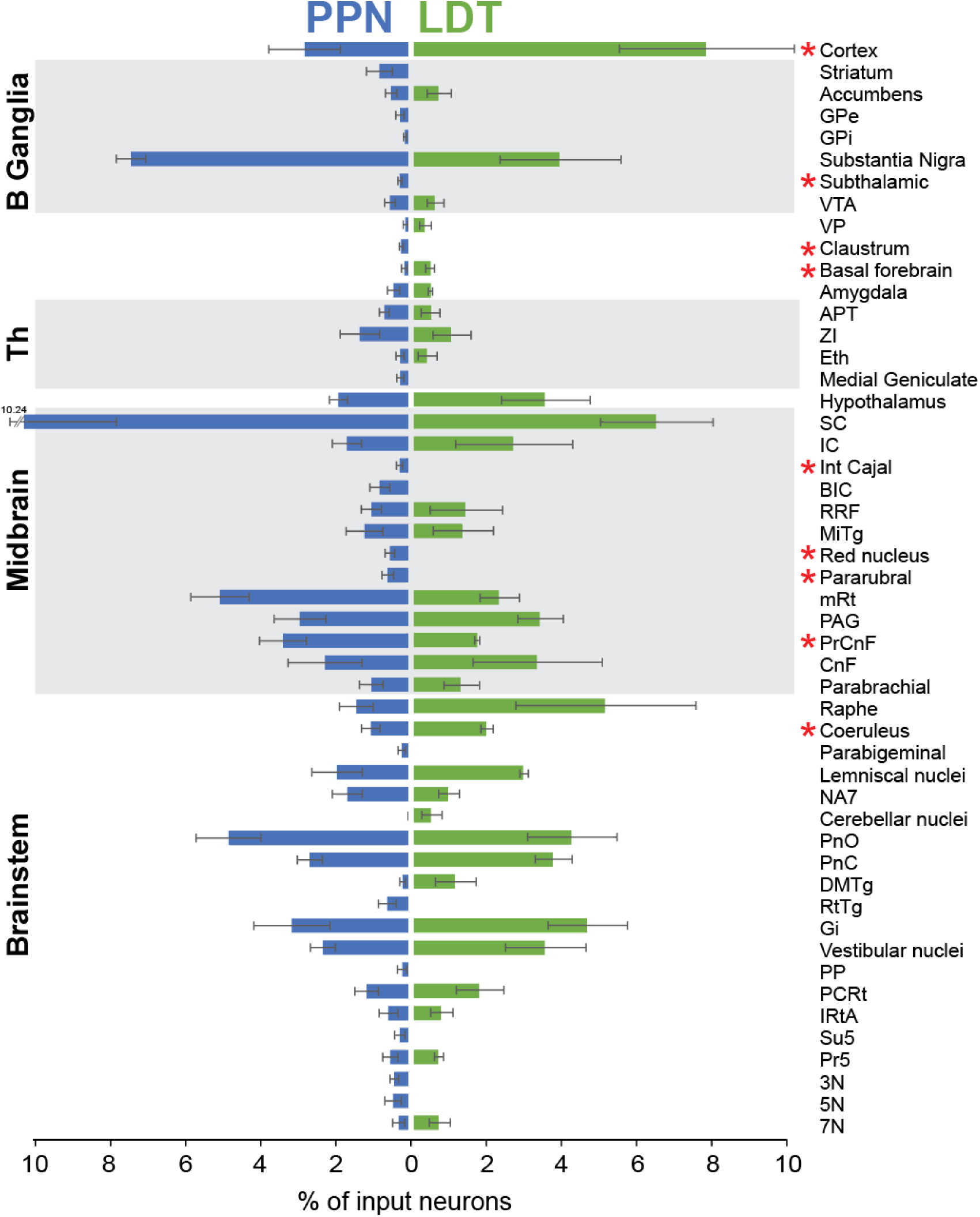
Major input structures to the cholinergic neurons of the PPN and LDT. Histogram showing the percentage of eGFP+ input (transsynaptically-labeled) neurons after injections of modified rabies virus in the PPN and LDT.The percentage is normalized by the total number of input neurons and the structures were organized in functional areas. Two-sided t-tests were used to compare differences between the number of input neurons to the PPN and LDT from all structures (*, P ≤ 0.05). Mean ± SEM (PPN n=6 rats; LDT n= 3 rats).

**Figure 4.**
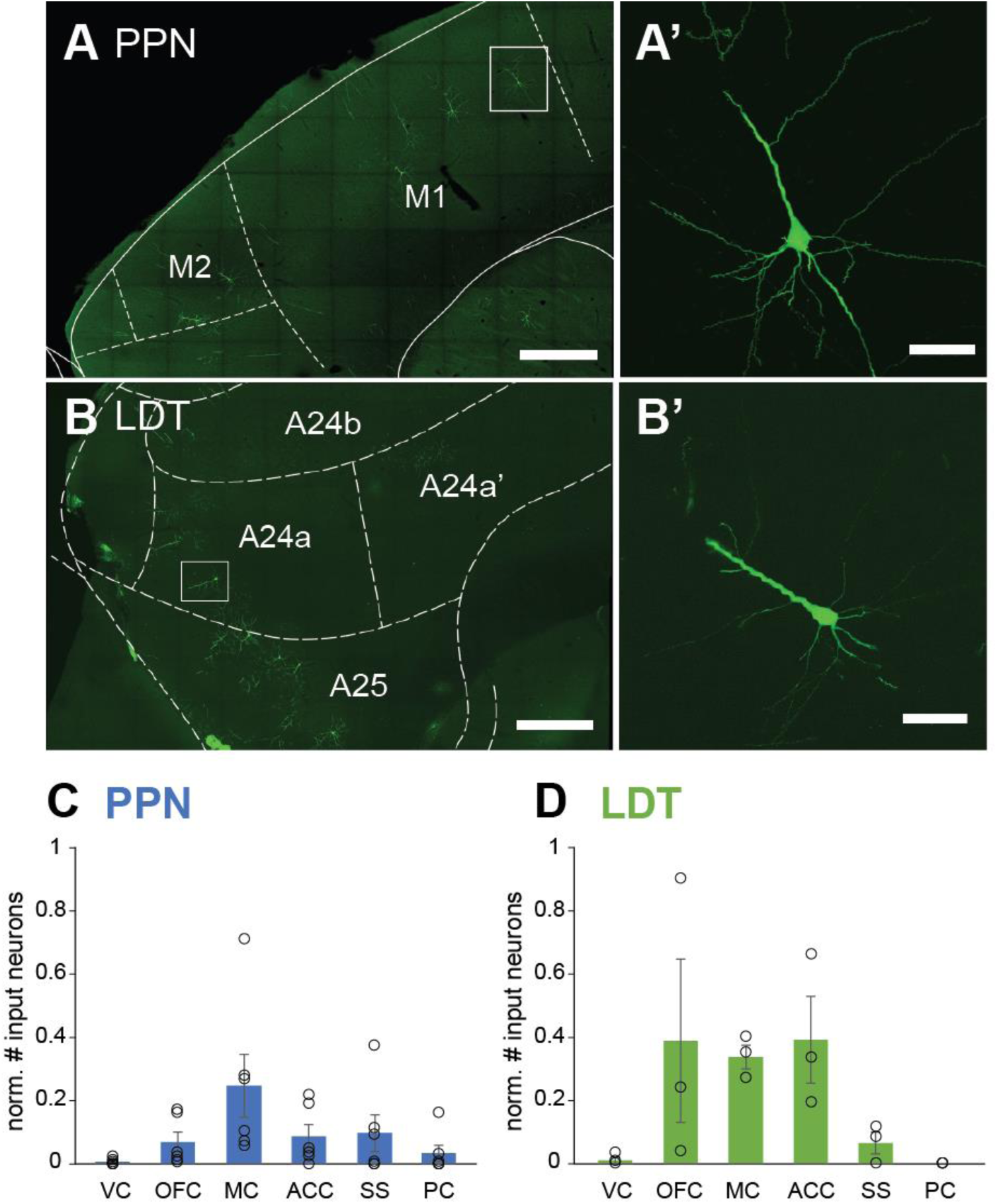
Distinct distribution of input neurons across cortical regions. **A**, Fluorescent micrograph showing cortical input neurons to PPN cholinergic cells located in the motor cortex in a sagittal section. **A’**, High magnification image of neuron in M1 (box in A). **B**, Fluorescent micrograph showing input neurons to the LDT in the anterior cingulate cortex in a sagittal section. **B’**, High magnification image showing a neuron located in A24a (box in B). **C**, Number of input neurons across cortical areas that innervate the cholinergic neurons of the PPN (**C**) and the LDT (**D**), normalized to the number of starter neurons. VC, visual cortex; OFC, orbitofrontal cortex; MC, motor cortex (M1/M2); ACC, anterior cingulate cortex; SS, somatosensory cortex; PC, parietal cortex. Mean ± SEM (PPN n=6 rats; LDT n=3 rats). Scale bars: A and B: 500 µm, A’ and B’: 100 µm.

**Figure 5.**
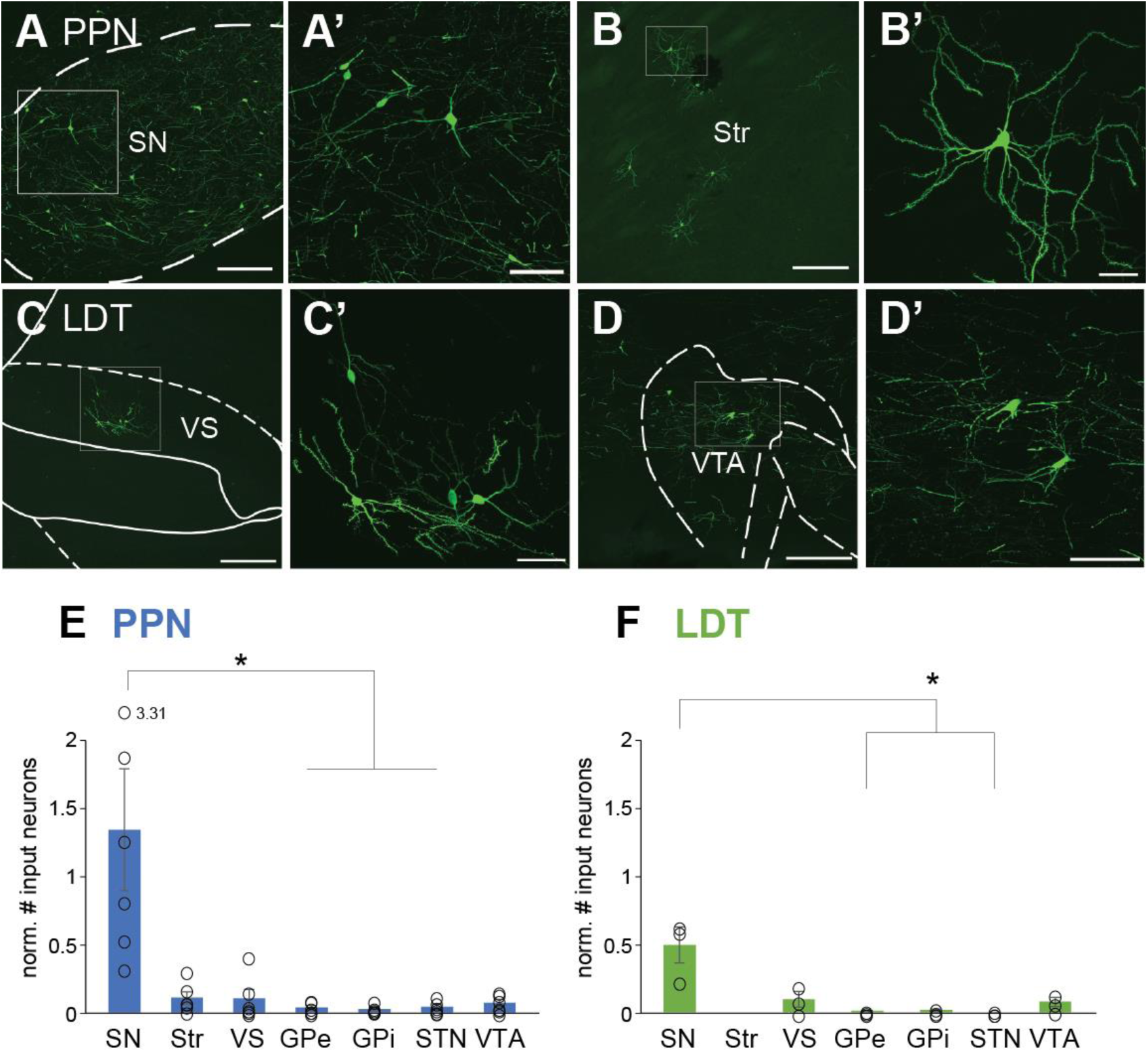
Basal ganglia input neurons predominantly target PPN. **A-D**, Fluorescent micrographs showing basal ganglia input neurons that target the PPN (**A-B**) and LDT (**C-D**). Neurons located in the substantia nigra (SN) were predominantly observed after PPN injections (**A-A’**). Input neurons in the dorsal striatum were only observed following PPN injections (**B-B’**); note the presence of multiple spines in the dendritic arbor of the labeled neuron, suggesting that it is spiny projection neuron. In contrast, LDT injections produced labeling of input neurons in the ventral striatum (VS; **C-C’**) and the ventral tegmental area (VTA; **D-D’**). The normalized number of input neurons (to the number of starter neurons) reveal a differential distribution between the PPN (**E**) and LDT (**F**). For both cases, the majority of neurons localize in the SN. Mean ± SEM (PPN n=6 rats; LDT n=3 rats). Kruskal-Wallis H test and Bonferroni corrected post hoc tests were performed to determine statistically significant differences, see text for values. Scale bars: A, B, C and D, 250 µm. A’ and D’ 100 µm, B’ and C’ 50 µm.

## Results

### Selective targeting of cholinergic neurons in midbrain nuclei

To characterize and map in whole-brain sections the neurons that innervate the cholinergic neurons of the PPN and the LDT, we used a monosynaptic retrograde tracing strategy (**Fig. 1A;** Callaway and Luo, 2015). For this purpose, we first injected into the PPN and the LDT of male and female ChAT::Cre rats (**Fig. 1B**) a combination of two helper viruses to induce the conditional (FLEX) expression of a TVA receptor and the rabies glycoprotein (G) in cholinergic neurons. This was followed 2 weeks later by an injection of G-deleted rabies virus in the same locations (SADΔ-eGFP; **Fig. 1A**). Neurons expressing the TVA receptor were tagged with mCherry (**Fig. 1C-D**) and neurons infected with the rabies virus were tagged with the enhanced green fluorescent protein (eGFP; **Fig. 1C’-D’**); neurons that were positive for both fluorescent reporters (mCherry+/eGFP+) were considered ‘starter neurons’ (**Fig. 1E-F**). Choline acetyltransferase (ChAT) immunolabeling was used to determine the cholinergic phenotype of starter neurons and locate the injection sites within the PPN/LDT rostro-caudal distribution (**Fig. 1C’’, 1D’’**). Injections in the PPN were targeted to the rostral (n=3 rats) and caudal (n=3 rats) portions of the nucleus, injections in the LDT were targeted at its center (n=3 rats; **Fig. 1B**). Following initial analysis, the data obtained from both regions of the PPN (i.e. rostral and caudal) were pooled together (n=6 rats) given the similarity in the mapping of the areas projecting to each PPN region and the number of presynaptic neurons counted.

Monosynaptically-labeled eGFP input neurons were distributed in a wide range of brain regions including the cortex, basal ganglia, forebrain, thalamus, hypothalamus, midbrain and upper and lower brainstem. eGFP labeling was intensely distributed along the neurons dendritic arbor and axonal projections (**Fig. 1E’**). Animals that had injections out of target, that had injections that overlapped between cholinergic nuclei, or where the quality of the labeling was poor, were not considered for further analysis. Control brains in which the helper viruses were omitted (n=3) did not display any type of presynaptic labeling, confirming the cell-type specificity for input tracing (not shown).

### Inputs to cholinergic neurons arise from widespread functional areas

To determine whether afferents to the distinct domains that compose the cholinergic midbrain originate in overlapping or separate brain regions, input neurons were initially grouped according to large functional divisions, i.e. cortex, basal ganglia, forebrain, thalamus, hypothalamus, midbrain and brainstem (**Fig. 2A**). We found that the midbrain and brainstem regions provide the largest number of input neurons to both PPN and LDT; both structures receive significantly more afferents from these functional areas than from the forebrain and the thalamus (two-way ANOVA; interaction: F(6,49)=0.918; P=0.490; PPN main effects: F(6,48)=11.713; P<0.001; Bonferroni corrected pairwise comparisons: midbrain vs forebrain/thalamus P<0.001, brainstem vs forebrain/thalamus P=0.001; LDT main effects: F(6,48)=5.239; P<0.001; Bonferroni corrected pairwise comparisons: midbrain vs forebrain P=0.011; midbrain vs thalamus P=0.005, brainstem vs forebrain P=0.002, brainstem vs thalamus P=0.005). In the PPN, only the basal ganglia showed a comparable number of input neurons to the midbrain/brainstem after rabies tracing (basal ganglia vs midbrain P=0.904; basal ganglia vs brainstem P=0.535). In contrast, the LDT receives a larger number of cortical inputs compared to the PPN (see below). Overall, while PPN and LDT share a similar distribution of input neurons across distinct brain regions (**Fig. 2B**), and this particularly evident in the midbrain and brainstem, the most notable differences were observed in the basal ganglia (for PPN) and the cortex (for LDT).

### Monosynaptic inputs from individual brain structures

To determine the differences in the innervation to PPN and LDT cholinergic neurons from individual structures among the above-defined functional regions, we examined the total number of neurons across the brain structures that fulfilled the criteria described in the Methods (total: 50 structures; **Fig. 3**). Analysis of the proportional contribution of each structure relative to the overall total number of input neurons to the PPN and the LDT for each animal revealed that several structures in the midbrain and the basal ganglia preferentially innervate the PPN. We observed that the vast majority of inputs to PPN cholinergic neurons originated in SC (10.24% ± 2.45). Within the SC, input neurons were preferentially located in the deep gray layer (DpG, 38.5% ± 4.6) but neurons also distributed notably in the deep (DpWh, 23.12% ±3.44) and intermediate white layers (InWh, 18.82% ± 4.5) as well as in the intermediate gray layer (InG, 16.86% ± 1.23). The SC is equally an important input source for the LDT (6.40% ± 1.49); no significant differences were detected between PPN and LDT. Within the SC, the majority of input neurons to the LDT were also localized preferentially in the DpG (39.58% ± 5.54). Other midbrain structures such as the interstitial nucleus of Cajal (int cajal), the brachium of the inferior colliculus (BIC), the red nucleus, the pararubral, the mesencephalic reticular formation (mRT) and the precuneiform (PrCnf), were found to have significantly more input neurons innervating the PPN than the LDT (two-tailed t-tests, int cajal P=0.036, BIC P=0.009, red nucleus P=0.036, pararubral nucleus P=0.016, mRT P=0.052, PrCnf P=0.044). After the SC, the second most important input region to the PPN was the SN (7.39% ± 0.39) which included neurons located in both, the pars compacta and pars reticulata. Compared to the PPN-injected animals, LDT-injected animals showed half of the input neurons in the SN (3.855% ± 1.6), although the difference was not significant (P=0.07). Similarly, the globus pallidus pars interna (GPi) and the subthalamic nucleus (STN) provide more input neurons to the PPN than the LDT (two-tailed t-tests, GPi P=0.086, STN P=0.01). In contrast, the largest source of inputs to the LDT was the cerebral cortex (7.73% ± 2.31) and this is in stark contrast to the PPN (2.76 ± 0.95; two-tailed t-test P=0.047). Another important input region to the LDT was the raphe (5.05% ± 2.38), which included the dorsal raphe (DR), with its ventral and dorsal segments, and the median and paramedian raphe nuclei. In addition, the locus coeruleus contained more input neurons for the LDT than for the PPN (two-tailed t-test P=0.044); the remaining brainstem structures examined did not show any preference for PPN or LDT. These results support a differential afferent balance between PPN and LDT according to the functional circuits in which they are embedded, where PPN cholinergic neurons preferentially receive inputs from structures involved in motor functions, whereas LDT cholinergic neurons receive a larger number of inputs from structures involved in limbic functions.

### Cortical inputs to the cholinergic midbrain

To further characterize the differences in the afferents from the cerebral cortex to the cholinergic midbrain, we identified the different cortical regions where input neurons were present, revealing different patterns between PPN and LDT. Cortical input neurons were distributed along the mediolateral axis and localized in the motor cortex (**Fig. 4A**) and somatosensory areas (SS), the orbitofrontal cortex (OFC) and anterior cingulate cortex (ACC; **Fig. 4B**). Input neurons in the motor cortex were similarly distributed between M1 and M2 areas, and because there were no differences in the number of input neurons between them, the data was pooled and presented as M1/M2 (MC, motor cortex). We observed that various primary somatosensory cortical areas project to the cholinergic midbrain; these neurons were located in the barrel cortex, the frontal lobe, and cortices that receive sensory inputs from the upper lip, jaw, trunk, shoulder and limbs. Input neurons within the OFC were distributed in the medial orbital area, orbital area, ventral orbital area, ventrolateral orbital area and agranular insular areas dorsal and ventral segments, and input neurons in ACC were mainly located in areas A24a, A24b, A25 and A32. The quantification of input neurons across these cortical areas revealed that PPN receives more inputs from MC than from other cortical regions (Kruskal-Wallis H test: χ2(5) = 13.456, P=0.019; Bonferroni corrected post hoc tests: MC vs visual cortex: P=0.015; **Fig. 4C**). In contrast, the LDT receives most of its inputs from the OFC, ACC and MC (Kruskal-Wallis H test: χ2(5) = 13.141, P=0.022; **Fig. 4D**). These results reveal that midbrain cholinergic neurons receive a specialized cortical input preferentially from either the MC, in the case of the PPN, or limbic cortices, in the case of the LDT.

### Basal ganglia inputs to the cholinergic midbrain

To further characterize the differences in the afferents from the basal ganglia to the PPN and LDT, we compared the number of input neurons innervating each midbrain structure. The vast majority of inputs from basal ganglia structures to the PPN arises from the SN (SN, 7.39% ± 0.39; **Fig. 5A,E**). Notably, we found input neurons in the striatum (**Fig. 5B**), which were identified as spiny projection neurons (0.76% ± 0.34; **Fig. 5B’**) with no specific distribution within striatal regions. Other basal ganglia inputs include the ventral striatum (0.45% ± 0.15), GPe (0.21% ± 0.11), GPi (0.08% ± 0.03), VTA (0.48% ± 0.14) and STN (0.22% ±. 0.05). The number of inputs neurons in the SN, however, was significantly larger than in other basal ganglia structures (Kruskal-Wallis H test, χ2(6) = 17.379, P=0.008; Bonferroni corrected post hoc tests: SN vs GPi: P=0.009; SN vs STN: P=0.024, SN vs GPe: P=0.021). LDT cholinergic neurons also received basal ganglia inputs, although this innervation was more restricted compared to that observed for the PPN. While the SN also provided the largest input to the LDT (3.85 ± 1.6; Kruskal-Wallis H test, χ2(6) = 14.719, P=0.023; Bonferroni corrected post hoc tests: SN vs STN: P=0.043; SN vs GPe: P=0.043), the rest of the basal ganglia inputs were distributed in ventral structures, such as the ventral striatum (0.64 ± 0.32; **Fig. 5C**) and the VTA (0.54 ± 0.22; **Fig. 5D**). No inputs to the LDT were observed to arise in the dorsal striatum or the STN. These results suggest that basal ganglia inputs to the cholinergic midbrain are functionally segregated, where dorsal basal ganglia structures innervate PPN cholinergic neurons, whereas ventral basal ganglia structures innervate LDT cholinergic neurons, in line with the functional segregation observed in cortical and midbrain areas. Notably, the SN seems to be a point of convergence of afferents to both cholinergic regions.

### Immunohistochemical characterization of input neurons

Given that an important number of SN and DR neurons synapse onto PPN and LDT cholinergic neurons, we aimed to characterized them neurochemically on the basis of the immunohistochemical expression of tyrosine hydroxylase (TH) or tryptophan hydroxylase (TPH), to identify dopaminergic and serotonergic neurons, respectively. In the SN, we found a large number of input neurons localized within the pars compacta and closely intermingled with TH+ neurons (**Fig. 6A**). However, in all cases input neurons were immunonegative for TH (**Fig. 6A’**). Similarly, in the DR, all input neurons were clearly identified within the borders set by the TPH+ labeling (**Fig. 6B**), but none of the eGFP+ neurons were immunopositive for TPH (**Fig. 6B’**). We then reasoned that it is possible that the rabies-mediated labeling interferes with the immunohistochemical detection on infected neurons (see also Dautan et al., 2020). To test this possibility, we then incubated input neurons sections with an antibody against the ubiquitous neuronal marker NeuN, which produces widespread non-selective nuclear labeling in the brain, to determine whether this protein can be detected in rabies-infected input neurons. Similar to the immunodetection of TH and TPH, we were unable to identify immunopositive signal for NeuN among input neurons (**Fig. 6C**), suggesting that rabies labeling interferes with detection of neuronal markers when using immunohistochemistry. Given the anatomical distribution of input neurons in the SN and DR, however, it is likely that both dopaminergic and serotonergic neurons directly innervate cholinergic neurons of the PPN and the LDT.

**Figure 6.**
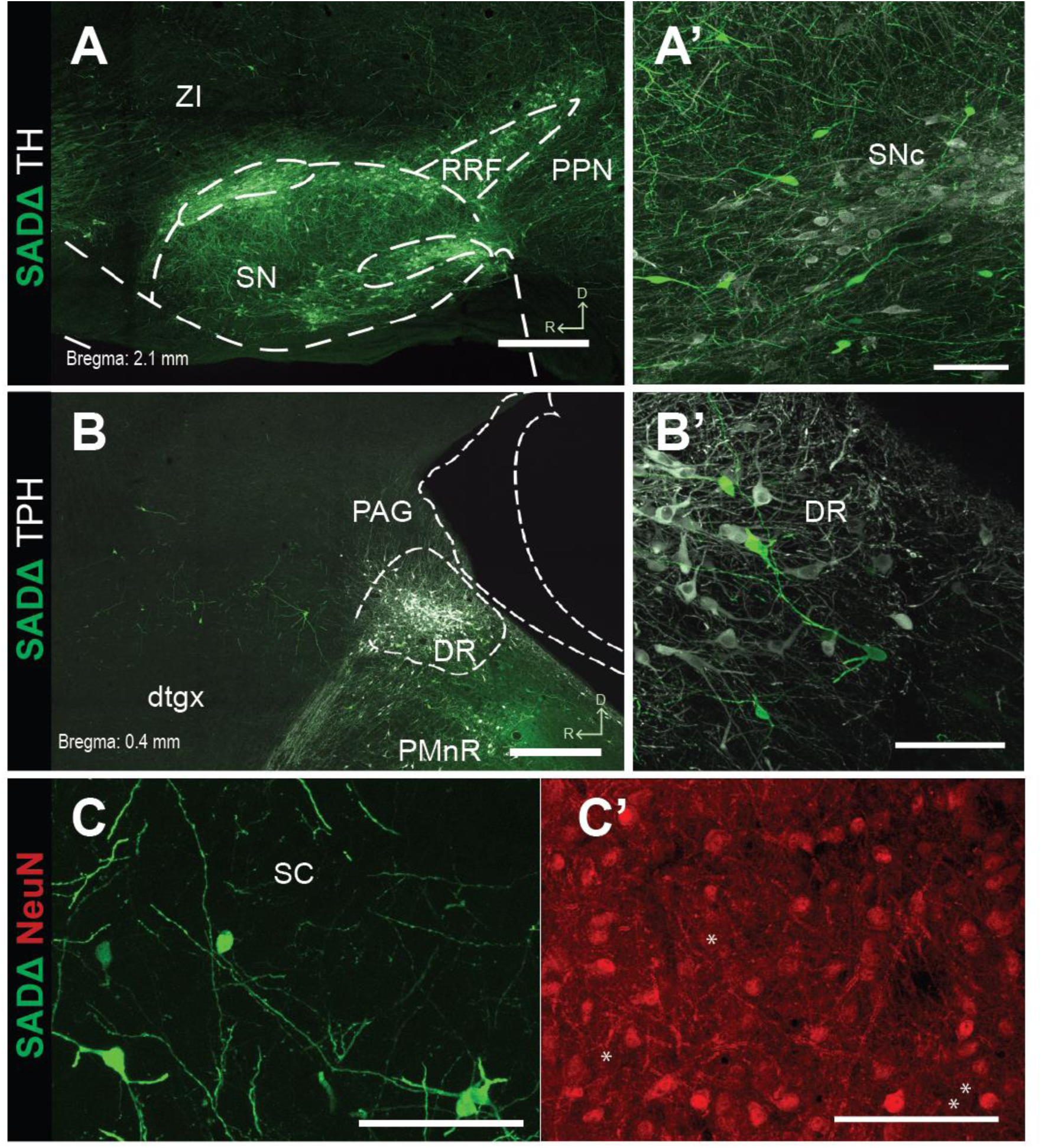
Neurochemical characterization of input neurons in the substantia nigra compacta and dorsal raphe. **A**, Input neurons located within the borders of the SNc after injections in the PPN and LDT were immunolabeled with a TH antibody to identify dopaminergic neurons. In all cases, input neurons were immunonegative, despite being closely intermingled with TH+ neurons (**A’**). **B**, Input neurons located within the borders of the DR after injections in the LDT were immunolabeled with a TPH antibody to identify serotonergic neurons. In all cases, input neurons were immunonegative, despite being closely intermingled with TPH+ neurons (**B’**). **C**, Input neurons located in the superior colliculus (SC) were incubated with NeuN antibodies to detect the presence of this ubiquitous neuronal protein. **C’**, SC input neurons were immunonegative for NeuN, thus confirming the limitations of immunohistochemical detection of neuronal markers in rabies-infected neurons. Scale bars: A &B 500 µm; A’, B’ & C 100 µm.

## Discussion

In the present study we used a retrograde transsynaptic strategy to characterize the identity and whole-brain distribution of afferents that selectively innervate midbrain cholinergic neurons. We show that these afferents are widely distributed across the brain and reveal fundamental differences with other brain cholinergic systems (e.g. basal forebrain and striatal cholinergic interneurons; see below). Given the established differences in output connectivity and function between PPN and LDT, we hypothesized that segregated afferent systems selectively synapse on each midbrain cholinergic subset. Our data show that PPN and LDT share some of the most prominent inputs (i.e. SC and SN), but notably differ in the functional segregation between motor and limbic afferents innervating PPN and LDT, respectively. Our results suggest that the cholinergic neurons of the midbrain operate as a heterogeneous functional entity that is capable of processing incoming motor and limbic signals which are in turn conveyed through parallel specialized circuits, an idea that was first introduced for basal ganglia-thalamocortical loops (Alexander et al., 1990), with which PPN/LDT maintain a close interconnectivity.

### Afferent overlap vs input selectivity

Our data show that the SC constitutes the main input structure to both the cholinergic PPN and LDT, in line with initial studies describing the connectivity between these structures (Semba and Fibiger, 1992). Input neurons in the SC were distributed in superficial, intermediate and deep layers, where there is a convergence of signals from visual, auditory and somatosensory modalities. Superficial layers receive input primarily from the retina and visual cortex. Deep layers, in contrast, receive inputs from several sensory modalities, inputs from motor areas, and projections from areas that are not purely sensory or motor (Cang et al., 2018). It is likely that such prominent input will transmit information about the orientation of the eyes, head and pinnae (Sparks, 1986), but also likely to signal target selection, attention (Krauzlis et al., 2013) and decision making (Wang et al., 2020).

In addition to the SC, we found that the SN also provides a prominent input to both the PPN and the LDT. It is well documented that the PPN maintains important reciprocal connectivity with most basal ganglia structures, but the position of the SN to modulate the LDT had not been established. Nigral projections arising from the pars reticulata have been described in detail (Beckstead et al., 1979; Edley and Graybiel, 1983) and are known to be the major input to the PPN. These synapses have been electrophysiologically characterized *in vitro* and their stimulation induces inhibition of PPN neurons (Noda and Oka, 1984; Scarnati et al., 1987), which suggests that this projection is largely GABAergic in nature (Childs and Gale, 1983). Interestingly, we found a number of input neurons in the dorsal and ventral striatum from PPN- and LDT-injected animals, respectively. These striatal neurons were distinguished as spiny projection neurons, suggesting that midbrain cholinergic neurons form part of the striatal output, as also seen for PPN glutamatergic neurons (Roseberry et al., 2016; Caggiano et al., 2018). Further studies should determine the relevance of this projection and the striatal output system to which it belongs (i.e. direct vs indirect pathway). Our data thus support that the cholinergic midbrain is a crucial link for relaying signals from virtually all basal ganglia structures, possibly influencing thalamic circuits that transmit feed-forward information to basal ganglia and cortical circuits.

We reveal that the cortex provides an important number of input neurons to the cholinergic midbrain, particularly to the LDT. Cortical projections to the LDT have been reported using conventional neuronal tracers, predominantly originating from the medial prefrontal and orbital cortices (Satoh and Fibiger, 1986; Terreberry and Neafsey, 1987; Cornwall et al., 1990; Takagishi and Chiba, 1991; Semba and Fibiger, 1992). Our data shows that input neurons in the OFC were found mainly in LDT-injected animals, but some were also present in the PPN group, thus likely providing a substrate for the involvement of the cholinergic midbrain in behavioral flexibility (Dalton et al., 2016; Amodeo et al., 2017; Izquierdo, 2017; Murray and Rudebeck, 2018). Also notably, we found input neurons in the anterior cingulate cortex mainly in the LDT group, which supports an involvement of the LDT in reinforcement learning and reward-guided selection of actions (Bohn et al., 2003; Cardinal et al., 2003; Schweimer and Hauber, 2005; Hillman and Bilkey, 2010). In contrast, we reveal a prominent input from MC (M1, M2) to the cholinergic PPN neurons. While the role of the cholinergic PPN in motor behavior is still elusive (see Introduction; also discussed in Gut and Mena-Segovia et al., 2019), a projection from the MC may reveal a role for PPN cholinergic neurons in movement preparation or a readiness-to-respond signal. Interestingly, however, the PPN has efferent connectivity with motor regions of the lower brainstem, such as the pontine nucleus part oral (PnO) and caudal (PnC), gigantocelullar nucleus (Gi) and spinal trigeminal nuclei, as well as projections to the spinal cord (Rye et al., 1988; Spann and Grofova, 1989; Grofova and Keane, 1991; Martinez-Gonzalez et al., 2014; Chen et al., 2017). This suggests that motor commands generated in cortical motor areas are transmitted to PPN cholinergic neurons and relayed to brainstem motor regions. Notably, we found a large number of input neurons in motor nuclei of the brainstem reticular formation, including the Gi, PnO, PnC and raphe magnus, suggesting a bidirectional role in the transmission of motor signals. Further experiments to fully elucidate the role of PPN cholinergic neurons in motor behavior are necessary.

### Input/output relationship of the cholinergic midbrain

Using conditional anterograde tracing of cholinergic neurons in ChAT::Cre rats, we have recently revealed the pattern of axonal innervation of PPN and LDT cholinergic neurons (Dautan et al., 2014, 2016a; Huerta-Ocampo et al., 2020). These previous studies together with our current dataset allow us to correlate the connectivity maps of midbrain cholinergic neurons. PPN cholinergic neurons maintain reciprocal connections with the basal ganglia, in particular the SN, GPe and striatum, the SC and motor nuclei of the brainstem, such as PnO, PnC and Gi. Interestingly, the connectivity with the cortex is also reciprocal, particularly with the motor and anterior cingulate cortices. In contrast, largely unidirectional efferent connectivity was observed with most thalamic nuclei, the ventral pallidum and amygdala. LDT cholinergic neurons maintain reciprocal connections with the inferior colliculus, DR, Gi, SN and ventral striatum. No LDT cholinergic axons were detected in the cortex, suggesting that this pathway is unidirectional. Similar to the PPN, we also found unidirectional efferent connectivity with the thalamus, the ventral pallidum and amygdala, but also with the globus pallidus, the olfactory tubercle and the medial and lateral septa. This comparative analysis based on our previously published results suggests that PPN and LDT maintain different levels of interconnectivity with the basal ganglia, the cortex and brainstem motor nuclei, whereas notably the thalamus stands out as a virtually exclusive output structure.

### Input specialization across brain cholinergic systems

Cholinergic neurons are distributed across the brain in anatomically defined cell clusters (Mesulam, 1990). Among them, cholinergic neurons of the basal forebrain and CINs share important functional properties with the cholinergic neurons of the PPN and LDT. For example, cholinergic neurons of the basal forebrain and the PPN/LDT increase their discharge during behavioral arousal (Jones, 2008). Moreover, CINs and cholinergic PPN/LDT neurons were found to be critical for the modulation of striatal output neurons during action selection (Dautan et al., 2020). These studies raise the question of whether cholinergic signaling from these systems is functionally overlapping. Recent studies have provided the identification of presynaptic inputs to the cholinergic basal forebrain (Do et al., 2016; Gielow and Zaborszky, 2017) and CINs (Guo et al., 2015; Klug et al., 2018). The comparison of the distribution of input neurons in these studies with the one reported here reveals that PPN/LDT cholinergic inputs are far more widely distributed across different brain regions than the inputs to any of these two other cholinergic groups. Basal forebrain neurons receive inputs predominantly from dorsal and ventral striatum, and from some cortical regions (orbital and insular) and the central nucleus of the amygdala. CINs receive inputs from several cortical regions, including M1, M2, S1, V1 and CC. Notably, CINs receive a prominent input from the thalamus and GPe. Thus, PPN/LDT cholinergic neurons share striatal inputs with the basal forebrain and cortical inputs with the CINs, although the proportions seem to differ greatly. This suggests that, despite the functional overlap across these cholinergic systems, they are largely segregated in their inputs. On the other hand, the convergence from some of these afferent regions (i.e. striatum and cortex) on distinct subsets of cholinergic neurons suggests the existence of an underlying mechanism capable of integrating cholinergic signaling across distant brain areas.

## Acknowledgements

We thank Yixin Tong and Selime Askit for their technical support in the acquisition of confocal images. This research was supported by NIH grant NS100824 (J.M.S.), a NARSAD Young Investigator Award (J.M.S.) and Rutgers University.

## References

Alexander GE, Crutcher MD, DeLong MR (1990) Basal ganglia-thalamocortical circuits: parallel substrates for motor, oculomotor,. Prog Brain Res 85:119–146.

Amodeo LR, McMurray MS, Roitman JD (2017) Orbitofrontal cortex reflects changes in response–outcome contingencies during probabilistic reversal learning. Neuroscience 345:27–37.

Beckstead RM, Domesick VB, Nauta WJ (1979) Efferent connections of the substantia nigra and ventral tegmental area in the rat. Brain Res 175:191–217.

Bohn I, Giertler C, Hauber W (2003) Orbital prefrontal cortex and guidance of instrumental behaviour in rats under reversal conditions. Behav Brain Res 143:49–56.

Bolam JP, Francis CM, Henderson Z (1991) Cholinergic input to dopaminergic neurons in the substantia nigra: A double immunocytochemical study. Neuroscience 41:483–494.

Caggiano V, Leiras R, Goñi-Erro H, Masini D, Bellardita C, Bouvier J, Caldeira V, Fisone G, Kiehn O (2018) Midbrain circuits that set locomotor speed and gait selection. Nature 553:455–460.

Callaway EM, Luo L (2015) Monosynaptic circuit tracing with glycoprotein-deleted rabies viruses. J Neurosci 35:8979–8985.

Cang J, Savier E, Barchini J, Liu X (2018) Visual Function, Organization, and Development of the Mouse Superior Colliculus. Annu Rev Vis Sci 4:239–262.

Cardinal RN, Parkinson JA, Marbini HD, Toner AJ, Bussey TJ, Robbins TW, Everitt BJ (2003) Role of the anterior cingulate cortex in the control over behavior by Pavlovian conditioned stimuli in rats. Behav Neurosci 117:566–587.

Chen T-W, Li N, Daie K, Svoboda K (2017) A Map of Anticipatory Activity in Mouse Motor Cortex. Neuron 94:866-879.e4.

Childs JA, Gale K (1983) Neurochemical evidence for a nigrotegmental GABAergic projection. Brain Res 258:109–114.

Clarke PBS, Hommer DW, Pert A, Skirboll LR (1987) Innervation of substantia nigra neurons by cholinergic afferents from pedunculopontine nucleus in the rat: neuroanatomical and electrophysiological evidence. Neuroscience 23:1011–1019.

Cornwall J, Cooper JD, Phillipson OT (1990) Afferent and efferent connections of the laterodorsal tegmental nucleus in the rat. Brain Res Bull 25:271–284.

Dalton GL, Wang NY, Phillips AG, Floresco SB (2016) Multifaceted Contributions by Different Regions of the Orbitofrontal and Medial Prefrontal Cortex to Probabilistic Reversal Learning. J Neurosci 36:1996–2006.

Dautan D, Hacioglu Bay H, Bolam JP, Gerdjikov T V., Mena-Segovia J (2016a) Extrinsic Sources of Cholinergic Innervation of the Striatal Complex: A Whole-Brain Mapping Analysis. Front Neuroanat 10:1.

Dautan D, Huerta-Ocampo I, Gut NK, Valencia M, Kondabolu K, Kim Y, Gerdjikov T V., Mena-Segovia J (2020) Cholinergic midbrain afferents modulate striatal circuits and shape encoding of action strategies. Nat Commun 11.

Dautan D, Huerta-Ocampo I, Witten IB, Deisseroth K, Bolam JP, Gerdjikov T, Mena-Segovia J (2014) A major external source of cholinergic innervation of the striatum and nucleus accumbens originates in the brainstem. J Neurosci 34:4509–4518.

Dautan D, Souza AS, Huerta-Ocampo I, Valencia M, Assous M, Witten IB, Deisseroth K, Tepper JM, Bolam JP, Gerdjikov T V., Mena-Segovia J (2016b) Segregated cholinergic transmission modulates dopamine neurons integrated in distinct functional circuits. Nat Neurosci 19:1025–1033.

Do JP, Xu M, Lee SH, Chang WC, Zhang S, Chung S, Yung TJ, Fan JL, Miyamichi K, Luo L, Dan Y (2016) Cell type-specific long-range connections of basal forebrain circuit. Elife 5.

Edley SM, Graybiel AM (1983) The afferent and efferent connections of the feline nucleus tegmenti pedunculopontinus, pars compacta. J Comp Neurol 217:187–215.

Gielow MR, Zaborszky L (2017) The Input-Output Relationship of the Cholinergic Basal Forebrain. Cell Rep 18:1817–1830.

Grofova I, Keane S (1991) Descending brainstem projections of the pedunculopontine tegmental nucleus in the rat. Anat Embryol (Berl) 184:275–290.

Guo Q, Wang D, He X, Feng Q, Lin R, Xu F, Fu L, Luo M (2015) Whole-brain mapping of inputs to projection neurons and cholinergic interneurons in the dorsal striatum. PLoS One 10.

Hillman KL, Bilkey DK (2010) Neurons in the rat anterior cingulate cortex dynamically encode cost-benefit in a spatial decision-making task. J Neurosci 30:7705–7713.

Huerta-Ocampo I, Hacioglu-Bay H, Dautan D, Mena-Segovia J (2020) Distribution of midbrain cholinergic axons in the thalamus. eNeuro 7.

Izquierdo A (2017) Functional Heterogeneity within Rat Orbitofrontal Cortex in Reward Learning and Decision Making. J Neurosci 37:10529–10540.

Jones BE (2008) Modulation of cortical activation and behavioral arousal by cholinergic and orexinergic systems. In: Annals of the New York Academy of Sciences, pp 26–34. Blackwell Publishing Inc.

Klug JR, Engelhardt MD, Cadman CN, Li H, Smith JB, Ayala S, Williams EW, Hoffman H, Jin X (2018) Differential inputs to striatal cholinergic and parvalbumin interneurons imply functional distinctions. Elife 7.

Krauzlis RJ, Lovejoy LP, Zénon A (2013) Superior Colliculus and Visual Spatial Attention. Annu Rev Neurosci 36:165–182.

Kroeger D, Ferrari LL, Petit G, Mahoney CE, Fuller PM, Arrigoni E, Scammell TE (2017) Cholinergic, glutamatergic, and GABAergic neurons of the pedunculopontine tegmental nucleus have distinct effects on sleep/wake behavior in mice. J Neurosci 37:1352–1366.

Martinez-Gonzalez C, van Andel J, Bolam JP, Mena-Segovia J (2014) Divergent motor projections from the pedunculopontine nucleus are differentially regulated in Parkinsonism. Brain Struct Funct 219:1451–1462.

Mena-Segovia J (2016) Structural and functional considerations of the cholinergic brainstem. J Neural Transm 123:731–736.

Mesulam MM (1990) Chapter 26 Human brain cholinergic pathways. Prog Brain Res 84:231–241.

Murray EA, Rudebeck PH (2018) Specializations for reward-guided decision-making in the primate ventral prefrontal cortex. Nat Rev Neurosci 19:404–417.

Noda T, Oka H (1984) Nigral inputs to the pedunculopontine region: intracellular analysis. Brain Res 322:332–336.

Parent M, Descarries L (2008) Acetylcholine innervation of the adult rat thalamus: Distribution and ultrastructural features in dorsolateral geniculate, parafascicular, and reticular thalamic nuclei. J Comp Neurol 511:678–691.

Paxinos G, Charles Watson (2007) The Rat Brain in Stereotaxic Coordinates Sixth Edition.

Roseberry TK, Lee AM, Lalive AL, Wilbrecht L, Bonci A, Kreitzer AC (2016) Cell-Type-Specific Control of Brainstem Locomotor Circuits by Basal Ganglia. Cell 164:526–537.

Rye DB, Lee HJ, Saper CB, Wainer BH (1988) Medullary and spinal efferents of the pedunculopontine tegmental nucleus and adjacent mesopontine tegmentum in the rat. J Comp Neurol 269:315–341.

Satoh K, Fibiger HC (1986) Cholinergic neurons of the laterodorsal tegmental nucleus: Efferent and afferent connections. J Comp Neurol 253:277–302.

Scarnati E, Proia A, Di Loreto S, Pacitti C (1987) The reciprocal electrophysiological influence between the nucleus tegmenti pedunculopontinus and the substantia nigra in normal and decorticated rats. Brain Res 423:116–124.

Schweimer J, Hauber W (2005) Involvement of the rat anterior cingulate cortex in control of instrumental responses guided by reward expectancy. Learn Mem 12:334–342.

Semba K, Fibiger HC (1992) Afferent connections of the laterodorsal and the pedunculopontine tegmental nuclei in the rat: A retro- and antero-grade transport and immunohistochemical study. J Comp Neurol 323:387–410.

Spann BM, Grofova I (1989) Origin of ascending and spinal pathways from the nucleus tegmenti pedunculopontinus in the rat. J Comp Neurol 283:13–27.

Sparks DL (1986) Translation of sensory signals into commands for control of saccacid eye movements: Role of primate superior colliculus. Physiol Rev 66:118–171.

Steriade M, Paré D, Parent A, Smith Y (1988) Projections of cholinergic and non-cholinergic neurons of the brainstem core to relay and associational thalamic nuclei in the cat and macaque monkey. Neuroscience 25:47–67.

Takagishi M, Chiba T (1991) Efferent projections of the infralimbic (area 25) region of the medial prefrontal cortex in the rat: an anterograde tracer PHA-L study. Brain Res 566:26–39.

Terreberry RR, Neafsey EJ (1987) The rat medial frontal cortex projects directly to autonomic regions of the brainstem. Brain Res Bull 19:639–649.

Wang L, McAlonan K, Goldstein S, Gerfen CR, Krauzlis RJ (2020) A causal role for mouse superior colliculus in visual perceptual decision-making. J Neurosci 40:3768–3782.

Watabe-Uchida M, Zhu L, Ogawa SK, Vamanrao A, Uchida N (2012) Whole-Brain Mapping of Direct Inputs to Midbrain Dopamine Neurons. Neuron 74:858–873.

Wickersham IR, Lyon DC, Barnard RJO, Mori T, Finke S, Conzelmann KK, Young JAT, Callaway EM (2007) Monosynaptic Restriction of Transsynaptic Tracing from Single, Genetically Targeted Neurons. Neuron 53:639–647.

Witten IB, Steinberg EE, Lee SY, Davidson TJ, Zalocusky KA, Brodsky M, Yizhar O, Cho SL, Gong S, Ramakrishnan C, Stuber GD, Tye KM, Janak PH, Deisseroth K (2011) Recombinase-driver rat lines: Tools, techniques, and optogenetic application to dopamine-mediated reinforcement. Neuron 72:721–733.

Woolf NJ, Butcher LL (1986) Cholinergic systems in the rat brain: III. Projections from the pontomesencephalic tegmentum to the thalamus, tectum, basal ganglia, and basal forebrain. Brain Res Bull 16:603–637.

Xiao C, Cho JR, Zhou C, Treweek JB, Chan K, McKinney SL, Yang B, Gradinaru V (2016) Cholinergic Mesopontine Signals Govern Locomotion and Reward through Dissociable Midbrain Pathways. Neuron 90:333–347.

